# Acute stress impairs target enhancement rather than distractor suppression in attention selection: Evidence from the N2pc and PD

**DOI:** 10.1101/2022.09.16.508346

**Authors:** Jingqing Nian, Run Yang, Jiao Xie, Yu Zhang, Yu Luo

## Abstract

**Background:** Studies have shown that acute stress significantly impacts the selection of emotional stimuli. However, the extent to which acute stress affects the cognitive mechanisms underlying target enhancement and distractor suppression when selecting emotionally neutral stimuli remains unclear.

**Methods:** We explored this issue using the Maastricht Acute Stress Test (MAST), a visual search task, and event-related potential recordings. Eighty healthy adults participanted in the experiment, which required them to search for a specific target while ignoring a color singleton distractor.

**Results:** The MAST successfully induced a stress response in the stress group, as indicated by the higher levels of salivary cortisol, state anxiety, negative emotion, as well as lower levels of positive emotion. Importantly, the stress group showed a significantly smaller N2pc in the lateral target with middle distractor displays than the control group. However, no significant differences in P_D_ were observed in the middle target with lateral distractor displays.

**Conclusions:** These results suggest that acute stress impairs target enhancement rather than distractor suppression during the selection of attention. This impairment may be due to impaired prefrontal cortex function under acute stress. The present research provides new insight into how acute stress affects attention selection.

**Trial registration:** This study has been submitted for registration with the Chinese Clinical Trial Registry (ChiCTR) and is currently under review (PID: 274121). The status will be updated as soon as the review process is completed.

## 1. Introduction

Acute stress is an individual’s physiological and psychological response to unpredictable and uncontrollable environmental stressors^[1]^, which has profoundly affects brain regions associated with the attentional selection of emotional stimuli^[2]^, such as prefrontal cortex, amygdala and hippocampus. This occurs through the activation of the hypothalamus-pituitary-adrenal (HPA) axis and the sympathetic nervous system (SNS), resulting in the release glucocorticoids and catecholamines^[3–5]^. For example, individuals have been shown to exhibit target enhancement^[6]^and distractor suppression impairments^[5]^for threat-related stimuli in a stressful context. Although acute stress is commonly studied in emotional contexts, it can also affect attentional selection in neutral situations. Research into anxiety suggests that group differences in attention are more pronounced under neutral conditions and have reduced effects in the presence of emotional content^[7,8]^. This highlights the need to examine the effects of stress on attention to neutral stimuli, as well as the cognitive mechanisms involved.

Attentional selection refers to the neural filtering processes that prioritize relevant stimuli over irrelevant distractors during the processing of visual information. This function is supported by two distinct mechanisms: target enhancement and distractor suppression. These mechanisms were originally proposed in early theoretical work by LaBerge^[9]^ and supported by ERP evidence from Hickey et al.^[10]^. Though these mechanisms are functionally and neurally distinct, they work together to resolve competition for limited attentional resources^[10–14]^. Target enhancement involves enhancing relevant features to facilitate target processing^[15]^ and requires the involvement of the frontal-parietal network^[16]^. In the ERP literature, target enhancement is indexed by the N2-posterior-contralateral (N2pc) component, which is a negative deflection observed over posterior electrodes contralateral to the attended item^[17–20]^. The N2pc is considered a neural marker of attentional selection that reflects the allocation of spatial attention to task-relevant stimuli. However, it can also be elicited by salient distractors that involuntarily capture attention^[21,22]^. Distractor suppression refers to the inhibition of irrelevant or potentially distracting visual stimuli to minimize their impact on attention^[23–26]^ and requires the involvement of visual cortex regions and/or the frontal-parietal network^[27–30]^. In contrast to the N2pc, the distractor positivity (P_D_) component is elicited by salient or otherwise potentially distracting irrelevant items^[10,31,32]^. Larger P_D_ amplitudes are typically observed when such distractors are successfully suppressed.

Previous studies have shown that acute stress impairs target processing by reducing the function of the fronto-parietal network^[33,34]^. For example, Sänger et al.^[34]^ found that acute stress impairs target processing, as evidenced by a diminished N2pc response. However, the impact of acute stress on distractor suppression is unclear. Some studies suggest that acute stress may attenuate distractor suppression when the suppression process involves the fronto-parietal network^[35,36]^. For instance, Xu et al.^[35]^ found that stress impaired distractor suppression in a visual search task, evidenced by a smaller P_D_ in the stressed group compared to the control group. Conversely, recent studies have found that the distractor suppression occurs in the visual cortex^[27,29,37,38]^, and a meta-analysis study revealed that acute stress does not impact neural activity in the visual cortex^[39]^. Therefore, if distractor suppression does not involve the fronto-parietal network, acute stress may not affect the distractor suppression.

Additionally, the effect of acute stress on attention selection may be mediated by distractor salience. Previous studies have shown that physically salient stimuli can capture attention in a bottom-up manner^[22,40–43]^. The Positivity posterior contralateral (Ppc) component may reflect the processing of the salience signals^[44–46]^. Furthermore, several studies have demonstrated that stressed individuals are more easily distracted by threatening (salient) stimuli that are task-irrelevant^[5,47]^. Gaspar and McDonald^[21]^found that the N2pc components appeared earlier in the high-anxiety group than in the low-anxiety group when a salient distractor appeared. This suggests that individuals with high anxiety may have difficulty suppressing distracting information, doing so more slowly. While previous studies have examined the effects of emotional or high-salience distractors on attention selection for neutral stimuli, the interaction between acute stress and the salience of a neutral distractor remains unclear. In short, this study investigated the cognitive mechanisms involved in how acute stress affects attention to neutral stimuli. Specifically, it examined whether acute stress impairs target enhancement and/or distractor suppression, as well as whether distractor salience modulates this effect. Participants completed a color-competition visual search task adapted from previous studies^[31,32]^. In this task, the spatial locations of the target and distractor were systematically manipulated to dissociate target enhancement from distractor suppression. This approach is based on well-established methods for isolating lateralized ERP components, such as the N2pc and P_D_. These components are elicited by simultaneous target and distractor singletons and serve as neural markers of distinct attentional processes^[10,22,44,48]^. Based on previous studies, we hypothesized that acute stress would impair target enhancement, resulting in a smaller N2pc in stressed participants than in non-stressed participants. However, we could not formulate a clear hypothesis regarding the effect of acute stress on distractor suppression because this process involves the visual cortex and/or the fronto-parietal network, which are affected differently by acute stress.

## 2. Method

### 2.1 Participants

Participants were recruited through advertisements on university campuses. To minimize the potential influence of sex hormones on cortisol measurements, all female participants were tested during the luteal phase of their menstrual cycles^[49]^. Prior to participation, participants were screened for normal color vision using the Ishihara color deficiency test. They were asked to confirm that they had no history of a diagnosed psychiatric disorders, habitual smoking, fainting, or epileptic seizures, and that they did not currently endocrine, autoimmune, cardiovascular, respiratory or dermatological disorders. They were also asked to confirm that they were not oversensitive to cutaneous sensation (in order to minimize negative reactions to cold water stress), that they were not using any medication that could affect cortisol response (e.g., β-blockers), and that they were not suffering from severe sleep disturbance or fatigue. Participants were asked to refrain from vigorous exercise, alcohol, and caffeine for three hours and to abstain from food for one hour, prior to the test. All participants were native speakers of Chinese.

The sample size was determined based on the method outlined in a previous studies^[21,32,35]^. A priori power analysis using G*Power (version 3.1) indicated that a sample size of 82 would provide over 80% power to detect a medium within-subject effect size (*f* = 0.314) at an alpha level of 0.05. A total of 87 participants were recruited. Seven participants were excluded from the final analysis due to more than 30 % of their EEG data trials being rejected. The final sample consisted of 80 healthy, right-handed participants (mean age: 20.95 ± 1.96 years; mean BMI: 20.97 ± 3.09; 45 female, 35 male).

The study protocol was approved by the Committee on Human Research Protection at the School of Psychology, Guizhou Normal University (protocol number 20190710), and was conducted in accordance with the Declaration of Helsinki. Participants provided written informed consent prior to the experiment and received 30 CNY per hour in compensation for their participation. The study was not pre-registered.

### 2.2 Experimental design

A full factorial 2×2 between-subjects design was used, with stress (stress, control) and distractor salience (high, low) serving as the manipulated between-subjects factors. Participants were randomly assigned to either the stress group (*n* = 41, 23 female) or the control group (no stress, *n* = 39, 22 female). After completing the Maastricht Acute Stress Test (MAST) procedure, participants were randomly assigned to either the high-salience distractor condition (stress: *n* =20,12 female; control: *n* =17, 11 female) or the low-salience distractor condition (stress: *n* =21, 11 female; ccontrol: *n* =22, 11 female).

### 2.3 General Procedure

The experiment was conducted between 13:30 and 18:30. Upon arrival, participants were escorted to the behavioral laboratory by an experimenter dressed either in a white lab coat and acting in a reserved manner (stress group), or in normal casual clothing and acting in a friendly manner (control group). As shown in Figure 1A, participants were then asked to rest for ten minutes. After this, they completed two questionnaires: the Positive and Negative Affect Schedule^[50]^ (PANAS) and the State-Trait Anxiety Inventory^[51]^(STAI-T; STAI-S). The first saliva sample (T1) was then collected using Salivette collection tubes (Sarstedt, Germany). Participants were asked to gently chew on a cotton swab to produce enough saliva. Participants then underwent EEG preparation while seated in a comfortable chair. This procedure lasted approximately 30 minutes. After the EEG set-up, the participants completed a second set of questionnaires and provided the second saliva sample (T2). Then, either the Maastricht Acute Stress Test (MAST) or a matched control task was performed for 15 minutes. Upon completion, a third set of questionnaires was completed and a third saliva sample (T3) was provided.

**Figure 1.**
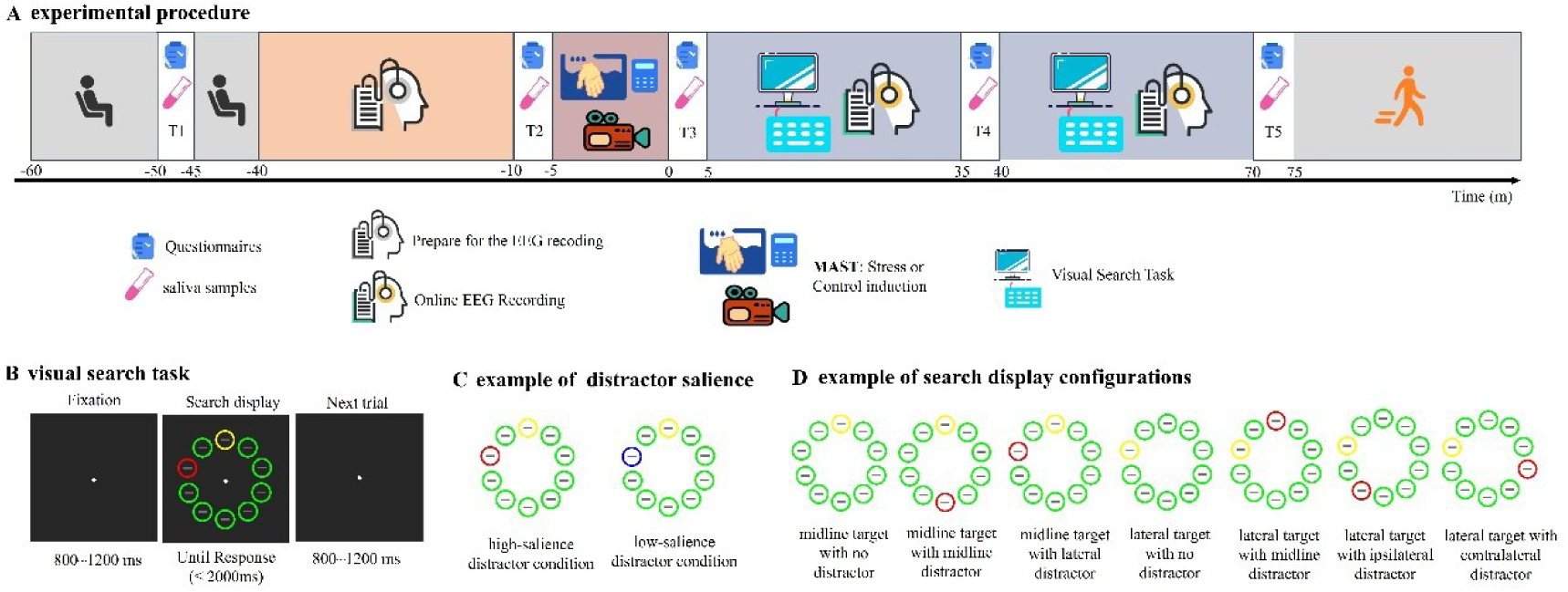
Overview of the experimental procedure and task. (A) Schematic illustration of the experimental procedure, (B) Example trial from the visual search task, (C) Examples of low- and high-salience distractor conditions, (D) Display configurations used across trials. In each trial, participants were instructed identify the orientation of the grey line inside the yellow target singleton. See the online article for the colour version of this figure.

Next, participants performed a visual search task while their EEG was being recorded. The visual search task consisted of four blocks. There was a 5-minute break between the second and third blocks. During this break, they completed a fourth set of questionnaires and provided a fourth saliva sample (T4). The visual search task took approximately 60 minutes to complete. After the EEG recording, participants completed a fifth set of questionnaires and provided a fifth saliva sample (T5). They were also asked to complete the Beck Depression Inventory (BDI) ^[52]^to assess their current mental health. Salivary cortisol samples were frozen at −20 °C within one hour of collection and stored until assayed. Cortisol was assayed by centrifugation at 3,000 rpm for 10 minutes to remove particulate matter. Salivary cortisol concentrations were determined using an electrochemiluminescence immunoassay (Cobas e 601, Roche Diagnostics, Nürbach, Germany), with a sensitivity threshold of 1.5 nmol/L. Following analysis, the salivary cortisol samples were destroyed as medical waste.

### 2.4 Stress induction

The Maastricht Acute Stress Test (MAST) was used to create either stressful or a non-stressul control condition^[53]^. The MAST is a well-established method of inducing stress. It involvs asking participants to immerse one hand in ice water while performing mental arithmetic in an unpredictable, uncontrollable environment involving social evaluation and negative feedback. Participants begad with a five-minutes preparation phase, followed by a ten-minutes acute stress phase. During this phase, participants sat in front of a computer and were briefed about the upcoming task. Those in the stress group, the participants were explicitly informed that their facial expressions would be recorded and that they would have to immerse their hand in ice-cold water (2.89 ± 0.32 °C, ranging from 1.9 to 3.4 °C) for a randomly selected period of 60–90 seconds, several times. Between each trial, they had to count backwards from 2043 in steps of 17 as quickly and accurately as possible. If they made a mistake, they received negative feedback from the experimenter and had to start again at 2043. Participants were instructed to continue with the mental arithmetic until the computer signalled the start of the next hand immersion trial, which took at least 45 s. The exact sequence and duration of the hand immersion trials (HIT) and mental arithmetic (MA) backward counting intervals are shown below: HIT-90 s, MA-45 s, HIT-60 s, MA-60 s, HIT-60 s, MA-90 s, HIT-90 s, MA-45 s, HIT-60 s. In the control condition, participants immersed their hands in water at body temperature (37.58 ± 0.51 °C, ranging from 36.5 to 38.6 °C) water and counted backwards from 2000 in steps of 10 at their own speed. No feedback was given and no facial expressions were recorded.

### 2.5 Visual Search Task

In the visual search task, the stimuli were presented on a 24.5-inch, 144-Hz AOC AG251FX LCD monitor using PsychToolbox-3 (http://psychtoolbox.org/). The monitor had a black background (RGB: 0, 0, 0) and was placed at a viewing distance of 57 cm.

The visual search task consisted of four blocks, with each participant completing a total of 1,260 trials. Each block contained 315 trials and a 5-second rest period was given after every 45 trials. After each block, participants were given a minimum 5-second rest period before starting the next block. Each participant was given at least 36 practice trials before the start of the experiment. During the breaks between blocks, participants received feedback on their compliance with the instructions, but not on their performance.

As shown in Figure 1B, each trial began with a central fixation for a random duration between 800 and 1,200 ms. This was followed by a search display consisting of ten unfilled circles, which were presented at equidistant around the central fixation at a distance of 9.2°. Each circle had a diameter of 3.4° and an 0.3° thick outline. A grey (RGB: 100, 100, 100) line oriented randomly in either the vertical or horizontal direction was inside each circle. Participants were instructed to fixation on the central point and identify the orientation (vertical or horizontal) of the grey line within the target singleton. They had to press one of two response buttons as quickly and accurately as possible to indicate this within 2,000 ms. The response buttons were counterbalanced across subjects. The next trial began after a response or no response within 2,000 ms.

In the visual search task, there were two types of trials: distractor-absent and distractor-present. In the former, nine circles were non-targets of a single color, and one circle was a yellow (RGB: 255, 255, 0) target. In the distractor-present trials, eight circles were non-targets of the same color, one was the yellow target singleton and one circle was a distractor singleton. As illustrated in Figure 1C, distractor salience was defined based on the chromatic contrast between the distractor and the surrounding non-targets. In the high-salience distractor condition, the distractor singleton (e.g., red; RGB: 255, 0, 0) was more distinct from the non-targets (e.g., green; RGB: 0, 255, 0) than the yellow target was from the background. In contrast, in the low-salience distractor condition, the distractor singleton (e.g., blue; RGB: 0, 0, 255) was less distinct from the non-targets (e.g., green) than the yellow target. Importantly, distractor salience was manipulated as a between-subjects variable to minimize participant fatigue and reduce the potential confounding effects of prolonged stress exposure.

To produce the visual search displays shown in Figure 1C, we varied the locations of the target and distractor. The following display configurations were produced: lateral target with no distractor (20.32%); midline target with no distractor (9.37%); lateral target with midline distractor (20.32%); lateral target with ipsilateral distractor (10.16%); lateral target with contralateral distractor (10.16%); midline target with lateral distractor (20.32%); midline target with midline distractor (9.37%). For the lateral configurations, 50% were on the left and 50% on the right. For the midline configurations, 50% were at the top and 50% at the bottom. These display configurations were then randomly intermixed across trials.

### 2.6 Electrophysiological Recording and Preprocessing

EEG recordings were made using a 64-channel electrode cap (Neuroscan, Herndon, VA, USA) according to the international 10/10 system with Ag/AgCl electrodes. The ground electrode was positioned between FPz and Fz, and the online reference electrode was placed between Cz and CPz. Eye movements and blinks were monitored using vertical electrooculograms (EOGs), which were recorded supra-orbitally and infra-orbitally relative to the left eye. The horizontal EOG was recorded as the difference in activity between the right and left orbital rim. The impedance of all electrodes was kept below 5 kΩ. EEG signals were recorded in DC mode using a Neuroscan SynAmps2 amplifier and Curry recording software. The signals were amplified with a low-pass filter at 100 Hz and digitised at a sampling rate of 1000 Hz, with no high-pass filters applied.

All analyses performed after data acquisition were carried out using the EEGLAB Toolbox^[54]^ and the ERPLAB Toolbox^[55]^. The EEG signals were referenced to the average of the left and right mastoids, and the four EOG signals were referenced using bipolar vertical and horizontal EOG derivations. All signals were then band-pass filtered using a non-causal Butterworth impulse response function with half-amplitude cutoffs at 0.1 and 30 Hz and a 12 dB/oct roll-off, before being resampled at 500 Hz. Portions of the EEG containing large muscle artefacts or extreme voltage offsets (identified by visual inspection) were removed. Additionally, segments were excluded from further analysis if the absolute voltage exceeded 100 μV per channel. Before independent component analysis (ICA), trials with extended inter-trial intervals (>2,000 ms between event markers) were excluded because they likely contained substantial ocular activity. ICA was then performed on the scalp EEG for each subject to identify and remove components that were associated with blinks^[56]^ and eye movements^[57]^. Previous studies have shown that artifact correction is reasonably effective at reducing these confounding factors, though it usually does not eliminate them entirely^[58]^. The ICA-corrected EEG data were then segmented for each trial from −100 to +500 ms relative to the onset of the search array. Baseline correction was based on the prestimulus time interval (−100 to 0 ms).

To minimize the impact of residual blinks or saccades on the EEG components in the epoch data, blinks were detected using a moving window peak-to-peak threshold (test period: 0-400 ms; moving windows full width: 200 ms; window step: 100 ms; voltage threshold: 80 μV) to identify an absolute amplitude of the vertical EOG exceeding 80 μV. Saccades were defined by using step-like artifacts (test period: 0-400 ms; moving windows full width: 200 ms; window step: 50 ms; voltage threshold: 40 μV) to detect a horizontal EOG amplitude difference greater than 40 μV. Trials containing incorrect responses, blinks, or saccades occurring between 0 and 400 ms were excluded from the analysis. This resulted in the retention rate of 93.58% for RT cutoffs and EEG artefacts. The data of seven participants who had more than 30% of their trials rejected due to artefacts, incorrect responses or response times (RTs) that were either too fast (RT < 300 ms) or too slow (RT > 1,500 ms) were excluded from the analyses.

### 2.7 Data Analysis

The data were analysed using JASP (version 0.16.3.0, 2022). JASP is a freely accessible, open-source statistical analysis package. We report any necessary corrections in the event of analysis-specific violations of assumptions.

The datasets have been made publicly available via the Science Data Bank (ScienceDB) and can be accessed at: http://cstr.cn/31253.11.sciencedb.02116.

#### 2.7.1 Stress response

To control for the potential impact of trait anxiety on stress-related outcomes, an analysis of variance (ANOVA) was performed on trait anxiety scores, with two factors: group (stress, control) and distractor salience (high, low).

To test for successful stress induction, three repeated-measures analyses of variance (ANOVAs) were performed on salivary cortisol and subjective affect ratings, with group (stress, control) and distractor salience (high, low) as between-subjects factors and time (T1 to T5) as a within-subjects factor. For ANOVAs involving more than two levels of a factor, the Greenhouse–Geisser correction was applied to address the violation of the sphericity assumption, and the corrected *p*-values are reported.

Two measures of cortisol output were quantified. First, the area under the cortisol curve relative to baseline (AUCg) was calculated as the total cortisol concentration over the duration of the experiment. Secondly, the cortisol concentration specifically related to the increase due to the stress induction (AUCi) was calculated by subtracting the baseline value from the AUCg value. All calculations followed the trapezoid integration method as recommended in a previous study ^[59]^.

To rule out the potential impact of stress induction on participants’ depressive symptoms, an analysis of variance (ANOVA) was performed on Beck Depression Inventory (BDI) scores, with two factors: group (stress, control) and distractor salience (high, low).

#### 2.7.2 Behavioral Performance

In order to examine the behavioral effects of stress and distractor salience, three indices were computed: the attentional capture effect, the target enhancement effect and the distractor suppression effect. The attentional capture effect was calculated as the difference in RTs between trials with and without a distractor, serving as a manipulation check for distractor salience, larger differences indicated stronger attentional capture of distractor. The target enhancement effect was defined as the difference in RT between lateral target with middle distractor trials and lateral target with no distractor trials. Smaller differences indicated stronger enhancement of target processing. The distractor suppression effect was calculated as the difference in RTs between midline target with lateral distractor trials and midline target with no distractor trials. Smaller values reflect more effective suppression of distractor interference.

Each of these effects was analyzed using a two-way ANOVA with two factors: group (stress, control) and distractor salience (high, low).

#### 2.7.3 ERP amplitude

As similar to those of previous studies^[21,31,32,35,44,60]^, the Ppc, P_D_ and N2pc were defined as contralateral minus ipsilateral difference waves at the PO7/PO8 electrodes. To test whether lateral target initially elicit N2pc in the lateral target/middle distractor displays, the target N2pc was computed within a time window of 200 –350 ms. To test whether the lateral distractor initially elicits Ppc and P_D_ in the midline target/lateral distractor trial, the Ppc was computed within a time window of 110–160 ms and the P_D_ within a time window of 250–500 ms.

An analysis of variance (ANOVA) was performed on the mean amplitude of the N2pc, Ppc and P_D_, with two factors: group (stress, control) and distractor salience (high, low). To further explore the effects of acute stress on target enhancement and distractor suppression, we analysed the correlation between the area under the curve (AUC) of cortisol and the mean amplitude of N2pc, Ppc and P_D_.

## 3. Results

### 3.1 Stress measurement

Trait anxiety: As shown in Figure 2E, there was no significant main effect of the group [*F*(1,76) = 0.15, *p* = 0.70] or distractor salience [*F*(1,76) = 0.37, *p* = 0.55], nor a significant interaction between group and distractor salience [*F*(1,76) = 1.48, *p* = 0.23]. The results showed no significant differences in trait anxiety scores across groups.

**Figure 2.**
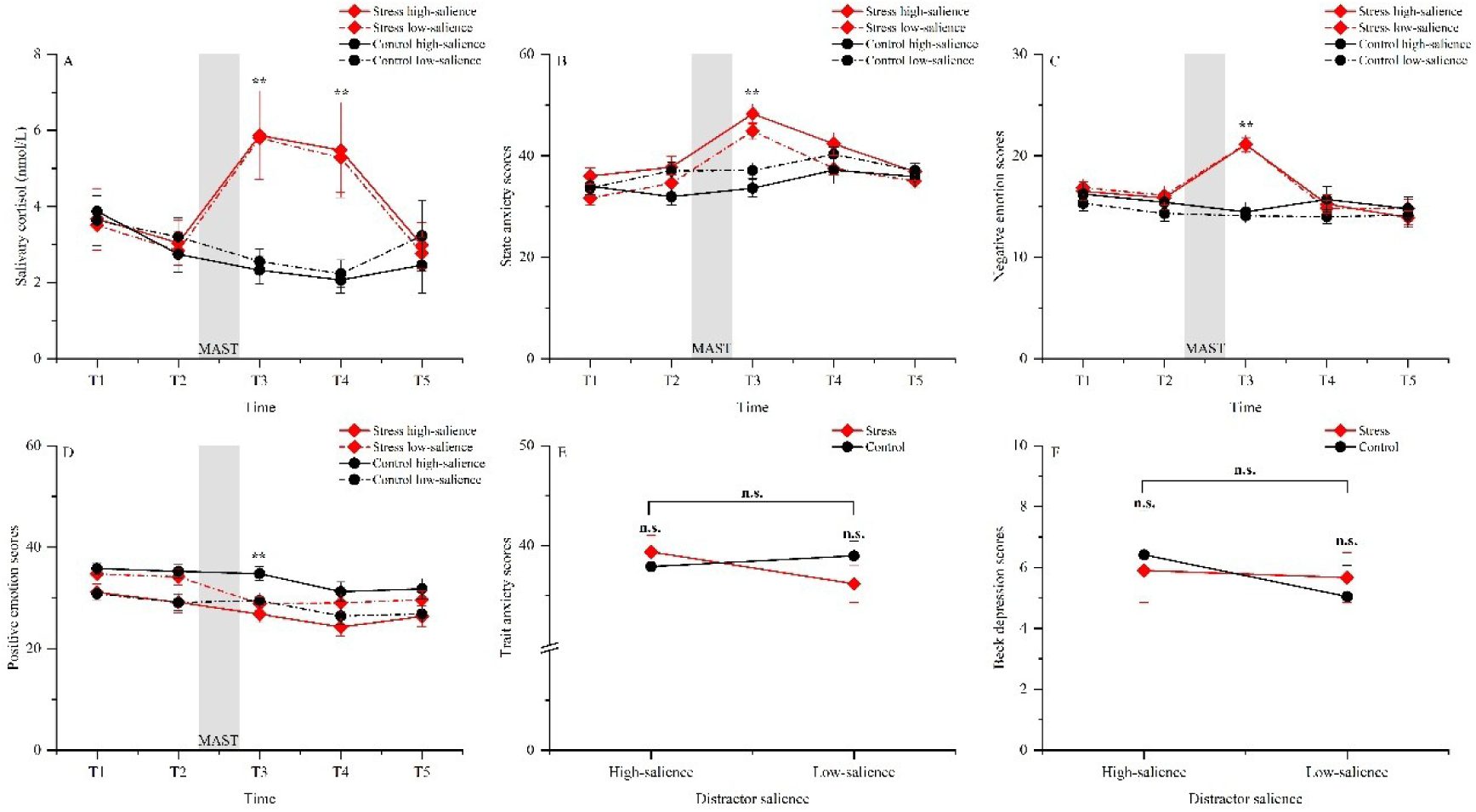
Results of stress response. Mean salivary cortisol levels (A), state anxiety scores (B), negative emotion scores (C), positive emotion scores (D), trait anxiety scores (E), and Beck Depression Inventory scores (F) for the stress and control groups across distractor salience conditions. At T3 (post-MAST), the stress group showed significantly higher cortisol, state anxiety, and negative emotion scores, and significantly lower positive emotion scores than the control group. Error bars represent standard errors of the mean. \*\**p*<0.01, \**p*<0.05, n.s. = not significant (*p* > 0.05). See the online article for the colour version of this figure.

Salivary cortisol: As shown in Figure 2A, the group main effect was significant [*F*(1,76) = 5.73, *p* = 0.02, *η^2^p* = 0.07]. The main effect of time was also significant [*F*(4,304) = 4.20, *p* = 0.003, *η^2^p* = 0.05]. The interaction between group and time was significant too [*F*(4,312) = 11.84, *p* < 0.01, *η^2^p* = 0.14]. Subsequent comparisons showed that the stress group had significantly higher cortisol levels than the control group at T3 [stress: 5.83 ± 4.50 nmol/L vs. control: 2.46 ± 1.48 nmol/L, *t* = 4.75, *p* < 0.01, *Cohen’s d* = 1.06] and at T4 [stress: 5.38 ± 4.85 nmol/L vs. control: 2.16 ± 1.57 nmol/L, *t* = 4.52, *p* < 0.01, *Cohen’s d* = 1.01]. No significant difference was observed between the two groups at the other time points [T1 and T2, *p*s > 0.05]. Within the stress group, a significantly higher cortisol levels were observed at T3 (5.83±4.50 nmol/L) and T4 (5.38 ± 4.85 nmol/L) compared to T2 (2.94 ± 2.43 nmol/L) and T1 (3.59 ± 3.22 nmol/L) [T3 v.s. T1, *t* = -4.30, *p* < 0.01, *Cohen’s d* = -0.71; T3 vs. T2, *t* = -5.56, *p* < 0.01, *Cohen’s d* = -0.91; T4 vs. T1, *t* = -3.42, *p* = 0.02, *Cohen’s d* = -0.56; T4 vs. T2, *t* = -4.68, *p* < 0.01, *Cohen’s d* =-0.77]. However, the main effect of distractor salience [*F*(1,76) = 0.01, *p* = 0.92], the distractor salience-time interaction [*F*(4,304) = 0.11, *p* = 0.98], the group-distractor salience interaction [*F*(1,76) = 0.17, *p* = 0.68], and the group-distractor salience-time interaction [*F*(4,304) = 0.14, *p* = 0.97] were not significant.

State anxiety: As shown in Figure 2B, the group main effect is marginally significant [*F*(1,76) = 3.65, *p* = 0.06, *η^2^p* = 0.05]. The main effect of time was also significant [*F*(4,304) = 29.90, *p* < 0.01, *η^2^p* = 0.28]. The interaction between group and time was significant too [*F*(4,304) = 20.28, *p* < 0.01, *η^2^p* = 0.21], Post-hoc analysis showed that the stress group had significantly higher state anxiety scores than the control group at T3 [stress: 46.51 ± 8.39 vs. control: 35.56 ± 7.16, *t* = 6.33, *p < 0.01, Cohen’s d* =1.42]. The interaction between group and distractor salience was also significant [*F*(1,76) = 4.48, *p* = 0.04, *η^2^p* = 0.06]. The post-hoc analysis revealed that the stress group exhibited significantly higher state anxiety scores than the control group in the presence of high distractor salience. However, the main effect of distractor salience [*F*(1,76) = 0.13, *p* = 0.72], the distractor salience-time interaction [*F*(4,304) = 1.31, *p* = 0.27], and the group-distractor salience-time interaction [*F*(4,304) = 1.23, *p* = 0.30] were not significant.

Negative emotion: As shown in Figure 2C, the main effect of group was significant [*F*(1,76) = 5.48, *p* = 0.02, *η^2^p* = 0.07]. The main effect of time was significant [*F*(4,304) = 18.18, *p* < 0.01, *η^2^p* = 0.19]. The interaction between group and time was also significant [*F*(4,304) = 22.62, *p* < 0.01, *η^2^p* = 0.23]. Post-hoc analysis showed that the stress group had significantly higher negative emotion score than the control group at T3 [stress: 21.12 ± 3.61 vs. control: 14.26 ± 3.63, *t* = 7.32, *p* < 0.01, *Cohen’s d* = 1.64]. However, the main effect of distractor salience [*F*(1,76) = 0.22, *p* = 0.64], the distractor salience-time interaction [*F*(4,304) = 0.55, *p* = 0.70], the group-distractor salience interaction [*F*(1,76) = 0.59, *p* = 0.45], and the group-distractor salience-time interaction [*F*(4,304) = 0.13, *p* = 0.97] were not significant.

Positive emotion: As shown in Figure 2D, the main effect of time was significant [*F*(4,304) = 34.69, *p* < 0.01, *η^2^p* = 0.31]. The interaction between group and time was significant too [*F*(4,304) = 4.46, *p* = 0.002, *η^2^p* = 0.06]. Post-hoc analysis showed that the stress group had significantly higher positive emotion scores than the control group at T3 [stress: 27.85±6.79 vs. control: 31.77±7.18, *t* = -2.54, *p* = 0.01, *Cohen’s d* = -0.57]. The interaction between group and distractor salience was also significant [*F*(1,76) = 8.46, *p* = 0.005, *η^2^p* = 0.10]. Post-hoc analysis revealed that the stress group exhibited significantly lower positive emotion scores than the control group in the presence of high distractor salience. However, the main effects of group[*F*(1,76) = 1.28, *p* = 0.26] and distractor salience [*F*(1,76) = 0.23, *p* = 0.63], the distractor salience-time interaction [*F*(4,304) = 0.59, *p* = 0.67], and the group-distractor salience-time interaction [*F*(4,304) = 0.91, *p* = 0.46] were not significant.

As a result, these salivary and subjective results suggest that the MAST was successfully used to induce the stress response in the stressed group.

Beck Depression Inventory: As shown in Figure 2F, There was no significant main effect of the group [*F*(1,76) = 0.003, *p* = 0.96] or distractor salience [*F*(1,76) = 0.55, *p* = 0.46], nor a significant interaction between group and distractor salience [*F*(1,76) = 0.28, *p* = 0.60]. The results showed no significant differences in trait anxiety scores across groups [Stress high-salience distractor: 5.90 ± 1.06 vs. Stress low-salience distractor: 5.67 ± 0.82 vs. Control high-salience distractor: 6.41 ± 1.43 vs. Control low-salience distractor: 5.05 ± 1.02].

### 3.2 Behavioral Results

Attentional capture effect: As shown in Figure 3A, neither the main effect of group [*F*(1,76) = 1.44, *p* = 0.23] nor the interaction between group and distractor salience [*F*(1,76) = 4.093×10^-6^, *p* = 0.99] was significant. However, the main effect of distractor salience was significant[*F*(1,76) = 14.40, *p* < 0.01, *η^2^p* = 0.16], indicating that the attentional capture effect was stronger for high-salience distractors than for low-salience ones.

**Figure 3.**
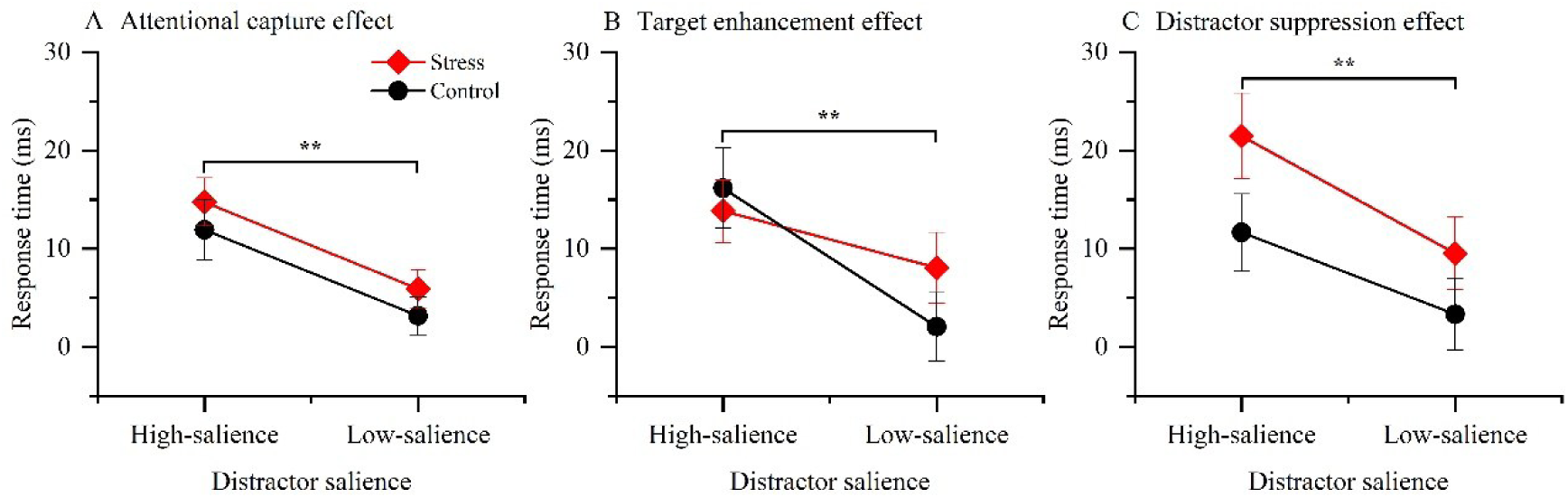
Behavioral results for the three computed effects. (A) The attentional capture effect is defined as the difference in response time between trials with and without a distractor. The high-salience distractor condition exhibited more attentional capture, though it was not affected by stress. (B) The target enhancement effect is calculated as the difference in response time between lateral target with midline distractor trials and lateral target with no distractor trials. The high-salience distractor condition had a lower target enhancement effect, which was not regulated by stress. (C) The distractor suppression effect, based on the response time difference between midline target with lateral distractor trials and midline target with no distractor trials. The distractor suppression effect was stronger for the control group than for the stress group, and it was stronger for low-salience distractors than for high-salience ones. \*\**p*<0.01, \**p*<0.05, n.s. = not significant (*p* > 0.05). See the online article for the color version of this figure.

Target enhancement effect: As shown in Figure 3B, neither the main effect of group [*F*(1,76) = 0.26, *p* = 0.61] nor the interaction between group and distractor salience [*F*(1,76) = 1.35, *p* = 0.25] was significant. However, the main effect of distractor salience was significant [*F*(1,76) = 7.64, *p* < 0.01, *η^2^p* = 0.09], indicating that the target enhancement effect was stronger for low-salience distractors than for high-salience ones.

Distractor suppression effect: As shown in Figure 3C, the main group effect was significant [*F*(1, 76) = 4.17, *p* = 0.045, *η^2^p* = 0.05], indicating that the distractor suppression effect was stronger for control group than for the stress group. The main effect of distractor salience was also significant [*F*(1,76) = 6.71, *p* = 0.01, *η^2^p* = 0.08], indicating that the distractor suppression effect was stronger for low-salience distractors than for high-salience ones. However, no significant interaction was found between group and distractor salience [*F*(1,76) = 0.21, *p* = 0.65].

### 3.3 Electrophysiological Results: ERP Components Target-elicited N2pc component

As shown in Figure 4, target-elicited N2pc component was calculated for lateral targets with midline distractor. The main group effect was significant [*F*(1, 76) = 5.06, *p* = 0.03, *η^2^p* = 0.06]. The main effect of distractor salience was marginal significant [*F*(1,76) = 3.14, *p* = 0.08, *η^2^p* = 0.04]. The interaction between group and distractor salience was significant [*F*(1,76) = 0.21, *p* = 0.65]. Post-hoc analysis revealed that the N2pc in the stress group (-1.07 ± 0.18 µV) was significantly smaller (more positive) than in the control group (-2.29 ± 0.37 µV) in the high-salience distractor condition [*t* = -3.31, *p* < 0.01, *Cohen’s d* = -1.09]. No significant results were observed in the low-salience condition (*p*s > 0.05).

**Figure 4.**
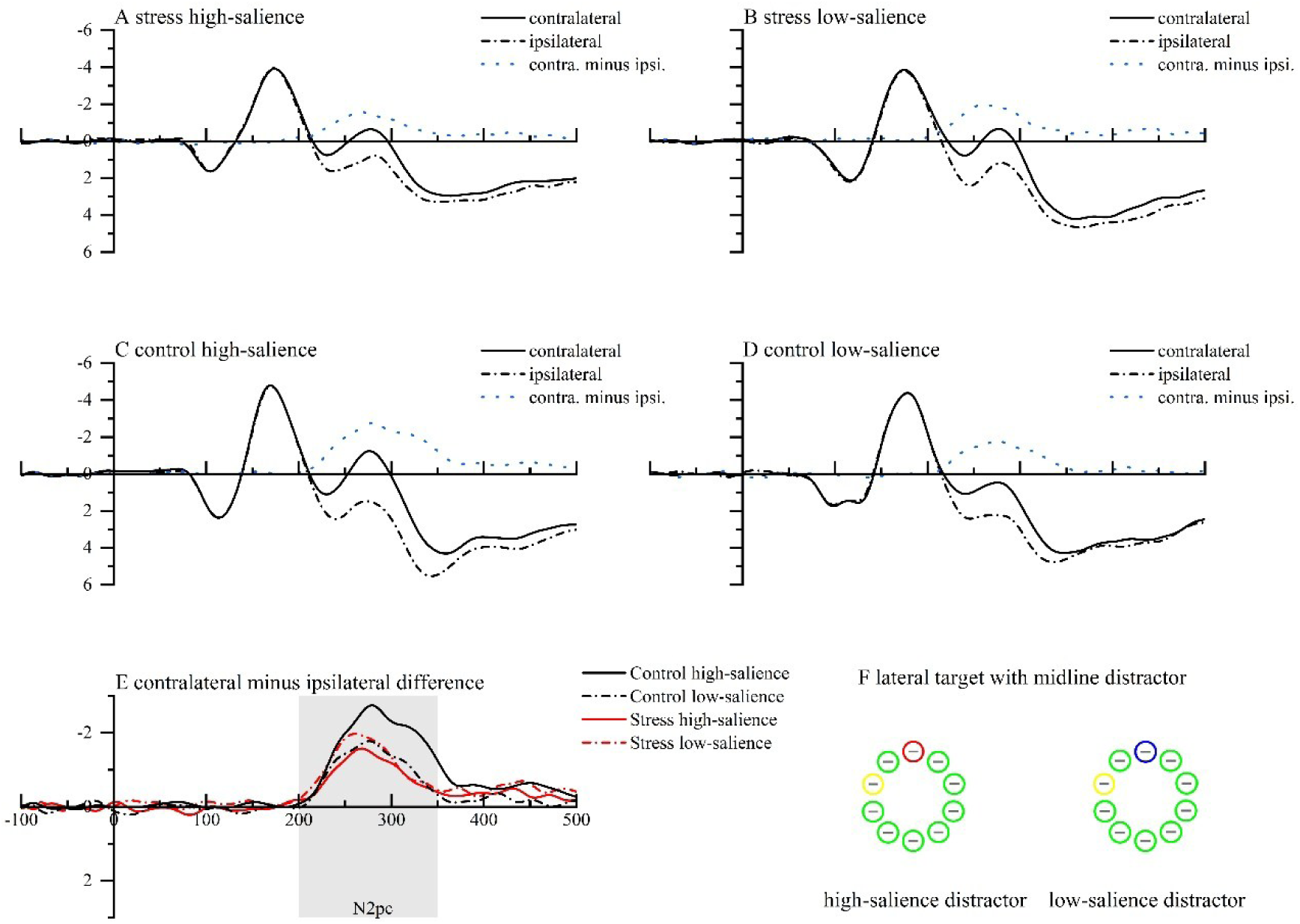
The ERP results for the lateral target with midline distractor across groups. Figures A–D show the grand-averaged ERPs recorded contralateral and ipsilateral to the target (with the distractor on the vertical midline), separately for each group and distractor salience. (E) The N2pc appears as the difference between the contralateral and ipsilateral waveforms. The shaded box represents the measurement windows for the target-elicited N2pc. Negative voltages are plotted upwards. See the online article for the color version of this figure.

#### Distractor-elicited Ppc and P_D_ components

**Ppc:** As shown in Figure 5, neither the main group effect [*F*(1,76) = 0.07, *p* = 0.79] nor the group-by-distractor salience interaction [*F*(1,76) = 1.12, *p* = 0.29] was significant. However, the main effect of distractor salience was significant[*F*(1,76) = 4.79, *p* = 0.03, *η^2^p* = 0.06]. The Ppc amplitude was larger in the high-salience distractor condition (0.48 ± 0.07 µV) than in the low-salience distractor condition (0.29 ± 0.06 µV).

**Figure 5.**
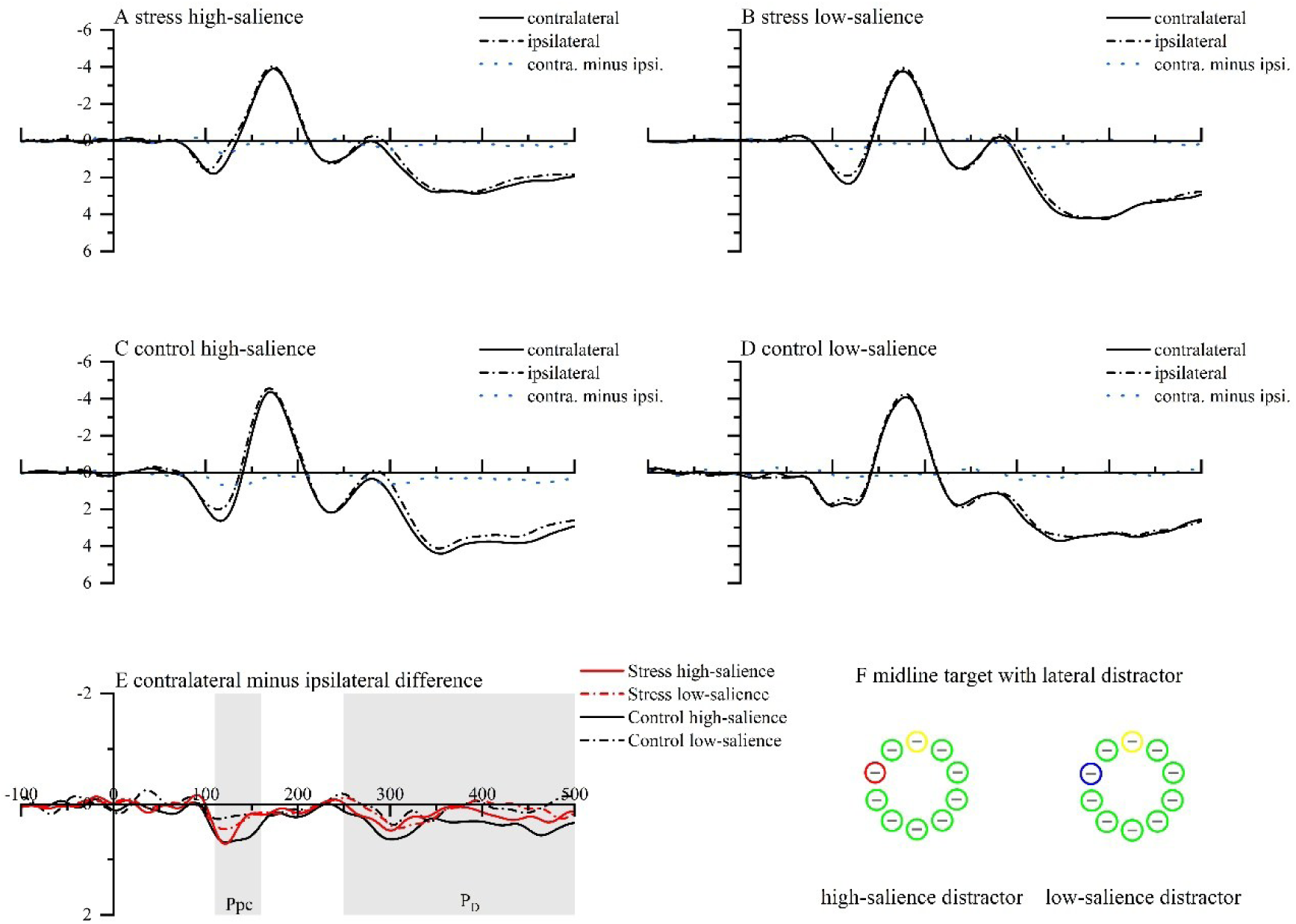
The ERP results for the midline target with lateral distractor across groups. Figures A–D show grand-averaged ERPs recorded contralateral and ipsilateral to the distractor (with the target on the vertical midline), separately for each group and distractor salience. (E) The Ppc and P_D_ appear as differences between the contralateral and ipsilateral waveforms. The shaded boxes represent the measurement windows for the distractor-elicited Ppc and P_D_. Negative voltages are plotted upwards. See the online article for the color version of this figure.

**P_D_:** As shown in Figure 5, neither the main effect of group [*F*(1,76) = 0.46, *p* = 0.50] nor the interaction between group and distractor salience [*F*(1,76) = 1.65, *p* = 0.20] was significant. However, the main effect of distractor salience was significant[*F*(1,76) = 4.42, *p* = 0.04, *η^2^p* = 0.06]. The P_D_ amplitude was larger in the high-salience distractor condition(0.29 ± 0.08 µV) than in the low-salience distractor condition(0.11 ± 0.05 µV).

### 3.4 Correlational Results

Spearman’s rank-order correlations revealed a negative correlation between the AUCi of cortisol and N2pc amplitude at the lateral target with midline distractor [*rs* = -0.24, *p* = 0.03]. However, no significant correlations were found between the AUCi of cortisol and Ppc [*rs* = -0.06, *p* = 0.63] or P_D_ [*rs* = -0.01, *p* = 0.90] amplitude at the midline target with lateral distractor.

## 4. Disscussion

The present study examined the effects of acute stress on target enhancement and distractor suppression when selecting attention for neutral stimuli. The MAST successfully induced a stress response, as indicated by higher levels of salivary cortisol, state anxiety, and negative emotions, as well as lower levels of positive emotions, in the stress group compared to the control group after MAST. In the high-salience distractor condition, the stress group showed a smaller N2pc than the control group. However, no significant difference was observed for the Ppc or P_D_. These results suggest that acute stress impairs target enhancement rather than distractor suppression during attention selection. This impairment may be attributed to the negative impact of acute stress on the PFC. These results offer valuable insight into the cognitive mechanisms through which acute stress affects attention to neutral stimuli.

We found that individuals responded more slowly in the high-salience distractor condition than in the low-salience distractor condition. The Ppc and P_D_ components were also more remarkable. These results suggest that the salience of the distractor was effectively altered, resulting in a delayed response to the target^[31,32,61]^. Additionally, we found that acute stress significantly reduced distraction suppression, though it did not affect attentional capture or target enhancement. While the stress group exhibited stronger interference effects under certain conditions, the interaction between group and salience was not significant. This finding suggests that the impact of stress on behavior is unstable. One study revealed that trait anxiety did not affect behavioral performance in a visual search task ^[21]^. This is because behavioral responses may result from multiple attentional processes acting together, making them difficult to attribute to a single factor, such as stress.

Importantly, we found that the N2pc amplitude was significantly smaller in the stress group than in the control group when high salience distractor was present. However, no group difference was observed for the P_D_ component. This pattern indicates that acute stress impairs target enhancement, which is consistent with findings in subclinical anxiety and depression^[62]^. Target enhancement is a PFC-dependent process^[63]^, whereas distractor suppression relies less on PFC functioning^[13]^. For example, Ort et al.^[63]^ found greater activation in the fronto-parietal control network when participants selected one of two potential targets among distractors compared to when only a single target was present. The observed impairment in target enhancement under stress is likely due to the detrimental effects of acute stress on PFC activity, which are well documented^[4,33]^. Similar findings have been reported with other stressors, such as sleep deprivation^[64]^, which disrupts attentional orienting as well. Consistent with this interpretation, our correlation analyses revealed that stronger activation of HPA was associated with greater impairment in target enhancement.

Furthermore, we found target enhancement impairments only under the condition of high-salience distractors. This result suggests that acute stress impairs selective attention under high-competition conditions, which is consistent with a previous study^[65]^. This selective impairment effect is most likely due to the PFC’s greater involvement in high-salience conditions than in low-salience conditions. Previous studies indicate that PFC involvement in top-down attentional control increases when task-irrelevant information (e.g., salience, task relevance, emotionality) can effectively compete for processing priority^[66]^. This suggests that the target processing in the high-salience distractor condition dependents heavily on the PFC and is thus more susceptible to stress ^[67]^. In contrast, less competition was involved when the distractor was not salient^[68,69]^. When competition is weak, less PFC is engaged, and target enhancement remains intact in acute stress situations^[70,71]^.

In contrast, we found no significant effect of acute stress on distractor suppression. This finding supports the idea that target facilitation and distractor suppression are governed by different cognitive processes^[11,10]^. Previous research has shown that the suppression of simple perceptual distractors (e.g., a red circle) occurs in the visual cortex and does not involve higher-order brain regions^[27,37,38,72,73]^, such as the PFC. Because the effects of acute stress are primarily mediated by the activation of the hypothalamic–pituitary–adrenal (HPA) axis, which targets the PFC^[3,4,74]^, it is reasonable to conclude that distractor suppression remains unaffected. This idea is further supported by a recent meta-analysis indicating that the visual cortex is unlikely to be involved in stress processing^[39]^. Thus, if the visual cortex is primarily responsible for distractor suppression, acute stress may have little or no impact on it. Further investigation of this hypothesis is warranted. However, this result contrasts with previous findings indicating that social stress impairs distractor suppression in visual search tasks^[35]^. A potential explanation for the inconsistent findings lies in the spatial gradient of distractor suppression. Previous studies have shown that distractors farther from the target are less likely to interfere, indicating ineffective attentional capture^[31]^. Xu et al.^[35]^ used displays with six items, resulting in shorter average distances between targets and distractors. This closer proximity likely amplified the effects of interference, making it easier to detect stress-related impairments in distractor suppression. In contrast, our study used ten-item displays. The increased spacing between targets and distractors, particularly in some trial configurations, may have reduced the strength of interference and thus masked the potential effects of acute stress. Future research should control target–distractor spatial relationships more precisely to better evaluate how acute stress influences distractor suppression.

Several limitations of the current study should be acknowledged. First, since the effects of acute stress last approximately 40 minutes^[53]^, the salience of the distractor was manipulated as a between-subjects factor to shorten the experimental duration. This design choice limited our ability to directly compare high- and low-salience conditions within the same individual. Future studies should use a within-subjects design to more precisely assess the interaction between stress and distractor salience. Second, the study may not have had sufficient power to detect subtle effects. Although the overall ANOVA revealed significant distractor salience effects, some pairwise comparisons, particularly those involving the Ppc and P_D_ components, were not significant, potentially due to ERP variability. To confirm these findings and better understand the effects of stress and salience on ERP components such as the Ppc and P_D_, future studies with larger samples or more trials are necessary.

## Conclusions

In conclusion, the current study shows that acute stress affects attention selection due to impairments in goal-directed target enhancement rather than difficulties in distractor suppression in the presence of a high-salience distractor. These impairments may be caused by the effects of acute stress on the PFC. The study provides clear evidence of acute stress-induced alterations in attention selection and reveals the cognitive mechanism by which acute stress affects attention selection.

## Abbreviations

PFC: prefrontal cortex
HPA: hypothalamus-pituitary-adrenal axis
SNS: sympathetic nervous system
MAST: Maastricht Acute Stress Test
Ppc: Positivity posterior contralateral
N2pc: N2 posterior contralateral
P_D_: Distractor Positivity
RT: response times

## Declarations

### Human Ethics and Consent to Participate declarations

The study protocol was approved by the Committee on Human Research Protection at the School of Psychology, Guizhou Normal University (Protocol:20190710) and conducted in accordance with the Declaration of Helsinki.

### Consent for publication

Not applicable.

### Availability of data and materials

The raw datasets used in these analyses have been made publicly available via the Science Data Bank (Science DB) and can be accessed at: http://cstr.cn/31253.11.sciencedb.02116.

### Authors’ contributions

JQN as lead for data curation, formal analysis and writing–original draft. RY served in a supporting role for data curation, formal analysis. JX served in a supporting role for data curation, formal analysis. YZ served as lead for conceptualization, supervision, writing–original draft, and writing–review and editing. YL served as lead for conceptualization, funding acquisition, supervision, writing–original draft, and writing–review and editing.

## Acknowledgements

Not applicable.

